# Transcutaneous vagus nerve stimulation reduces total striatal GABA content and facilitates early-phase motor learning

**DOI:** 10.1101/2025.08.08.669433

**Authors:** Kana Matsumura, Hiroyuki Matsuta, Ryushin Kawasoe, Tomoyuki Fumuro, Kojiro Matsushita, Nobuhiro Hata, Yoshiki Asayama, Tsuyoshi Shimomura, Minoru Fujiki, Hisato Sugata

**Affiliations:** Graduate School of Welfare and Health Science, Oita University, 700, Dannoharu, Oita 870-1192, Japan; Faculty of Medicine, Hospital Informatic Center, Oita University, 1-1, Idaigaoka, Hasama-machi, Yufu, Oita 879-5593, Japan; Department of Neurosurgery, School of Medicine, Oita University, 1-1, Idaigaoka, Hasama-machi, Yufu, Oita 879-5593, Japan; Division of Health Sciences, Graduate School of Medicine, Osaka University, 1-2, Yamadaoka, Suita, Osaka 565-0871, Japan; Department of Advanced Medical Sciences, Faculty of Medicine, Oita University, 1-1, Idaigaoka, Hasama-machi, Yufu, Oita 879-5593, Japan; Department of Mechanical Engineering, Gifu University, 1-1, Yanagito, Gifu 501-1193, Japan; Department of Radiology, Faculty of Medicine, Oita University, 1-1, Idaigaoka, Hasama-machi, Yufu, Oita 879-5593, Japan; Faculty of Welfare and Health Science, Oita University, 700, Dannoharu, Oita 870-1192, Japan; Graduate School of Medicine, Oita University, 1-1, Idaigaoka, Hasama-machi, Yufu, Oita 879-5593, Japan

**Keywords:** tVNS, gamma-aminobutyric acid, motor learning, magnetic resonance spectroscopy

## Abstract

**Background:** Transcutaneous vagus nerve stimulation (tVNS) has emerged as a promising non-invasive technique for modulating neuroplasticity. Previous studies have suggested that changes in regional brain GABA signaling contribute to these effects, but empirical neurophysiological evidence remains limited.

**Methods:** We investigated the neurophysiological and behavioral effects of tVNS (200-μs pulses at 20 Hz, alternating 30 s ON–1 s OFF cycles, 30 min total duration) in healthy adults using two experimental paradigms. In Experiment 1, GABA levels were measured in the left striatum (STR), dorsolateral prefrontal cortex (DLPFC), and sensorimotor cortex (SM) of 34 participants by magnetic resonance spectroscopy (MRS) before and after ipsilateral tVNS. In Experiment 2, 28 participants performed a right-hand force-control motor learning task before, during, and after tVNS.

**Results:** Administration of tVNS significantly reduced GABA levels in the left STR compared to sham stimulation (*p* < 0.05), and also significantly improved motor task performance compared to the sham group at 10 minutes after stimulus onset (*p* < 0.05)

**Conclusion:** Transcutaneous VNS may facilitate early-phase motor learning by reducing striatal GABA levels and consequently inducing corticobasal circuit disinhibition. These findings support tVNS as a potential noninvasive intervention to enhance motor learning for neurorehabilitation and motor disorder treatment.

## 1. Introduction

Invasive vagus nerve stimulation (iVNS) has been approved by the U.S. Food and Drug Administration as a treatment for epilepsies and depression as well as for stroke rehabilitation, and numerous clinical studies and reviews have confirmed the efficacy of this intervention [1,2]. For instance, a comprehensive review reported that iVNS reduced seizure frequency by 50%–100% in 45%–65% of patients [3]. However, iVNS requires surgical electrode implantation, and thus carries the risks of surgical side effects. Implantation surgery is also costly and not widely accessible, especially in developing countries. Conversely, transcutaneous vagus nerve stimulation (tVNS) provides a non-invasive, lower cost, and more accessible alterative [4–6], and in the two decades since it was first introduced has been reported to enhance social behavior in epilepsy patients [7–8], aid in post-stroke motor recovery [8–11], improve both motor and non-motor symptoms of Parkinson’s disease [10–12], provide symptomatic relief from depression [11–14] and post-traumatic stress disorder (PTSD) [13–15], and even suppress postoperative pain following total knee arthroplasty [14–16].

The primary acute outcome of tVNS is a shift in autonomic nervous system function toward parasympathetic dominance [7,8,15,16], but the downstream neurological changes underlying many of the reported therapeutic effects remain unknown. Recent evidence suggests that tVNS promotes neuroplasticity in midbrain and corticobasal pathways, potentially by modulating neurotransmitter metabolism and signaling [17]. For instance, tVNS via the left cymba conchae has been shown to activate the nucleus tractus solitarii and the locus coeruleus, leading to increased noradrenergic neurotransmission [18–22]. Another potential action mechanism is altered gamma-aminobutyric acid (GABA) signaling. GABA is the primary inhibitory neurotransmitter in the brain and several studies have reported the iVNS can increase neural GABA levels [23–26]. Also, long-term iVNS was reported to increase hippocampal GABA receptor density in the hippocampus of patients with drug-resistant epilepsy [26,27]. Thus, tVNS may exert therapeutic effects by modulating GABAergic signaling. In fact, this notion is indirectly supported by neuroimaging studies. Multiple studies have reported an inverse correlation between GABA levels and blood oxygen level-dependent (BOLD) signals [28–30]. For example, Stagg et al. [31] reported that decreases in GABA levels were associated with elevated BOLD signals in the sensorimotor cortex (SM), and Dolfen et al. [32] reported that increased striatal GABA was associated with reduced BOLD signals. Furthermore, Frangos and Komisaruk reported that tVNS increased BOLD signals in the basal ganglia, including the STR [33], suggesting a reduction in striatal GABA levels. Although there is growing interest in the neuromodulatory effects of tVNS for therapeutic intervention, the underlying mechanisms must be clarified. In particular, there is little direct neurophysiological evidence for modulation of GABAergic signaling by tVNS [25,34].

One especially promising application of tVNS is in motor rehabilitation, a therapeutic process that depends strongly on motor learning. Motor learning refers to the process by which individuals acquire new skills or refine previously learned skills, allowing them to adapt more effectively to an ever-changing environment [35,36]. This phenomenon involves structural and functional changes in several brain regions, such as STR [37], SM [38], and the dorsolateral prefrontal cortex (DLPFC) [38,39], and is associated with various regional changes in neurotransmitter signaling. Inhibitory control of corticobasal circuits mediated by GABAergic signaling is particularly important for motor learning [11,40], and several studies have reported that tVNS can enhance motor sequence learning [24,39] and promote memory consolidation skills in healthy individuals [41]. Reduced GABAergic activity is further implicated in these effects as tVNS has been shown to enhance BOLD activity, which in turn is negatively correlated with GABAergic transmission. However, several studies have reported that tVNS can also increase regional GABA [24,25,42,43]. Therefore, additional studies investigating the effects of tVNS on both GABA-associated neuroimaging signals and motor learning are required.

We speculated that tVNS promotes motor task performance by modulating GABA levels or signaling within STR, DLPFC, and (or) SM circuits. To test this hypothesis, we measured the effects of tVNS on GABA levels in the STR, DLPFC, and SM using proton magnetic resonance spectroscopy (^1^H-MRS) (Experiment 1), as all of these regions are implicated in motor learning, and further examined the effects of tVNS on the acquisition of a motor learning task (Experiment 2).

## 2. Materials and Methods

### 2.1. Ethics statement

All experimental protocols were approved by the Ethics Review Board of the Oita University Faculty of Welfare and Health Science (No. F240033) and conducted in accordance with the tenets of the Declaration of Helsinki. The study purpose and possible consequences were fully explained prior to experiments, and informed consent was obtained from all participants.

### 2.2. Transcutaneous vagus nerve stimulation

In both Experiments 1 and 2, the vagus nerve was stimulated noninvasively using a specialized tVNS device (Soterix Medical, Inc., USA). In the tVNS group, the stimulation electrode was placed on the left cymba conchae to target the auricular branch of the left vagus nerve, while in the corresponding sham group, electrodes were positioned on the left earlobe, an area not innervated by the vagus nerve and thus not expected to induce vagal activation [44–46] (Figure 1a). The tVNS device was set to deliver electrical stimulation with a pulse width of 200 μs at 20 Hz, alternating between 30 seconds ON and 1 s OFF, for a total duration of 30 minutes.

**Figure 1.**
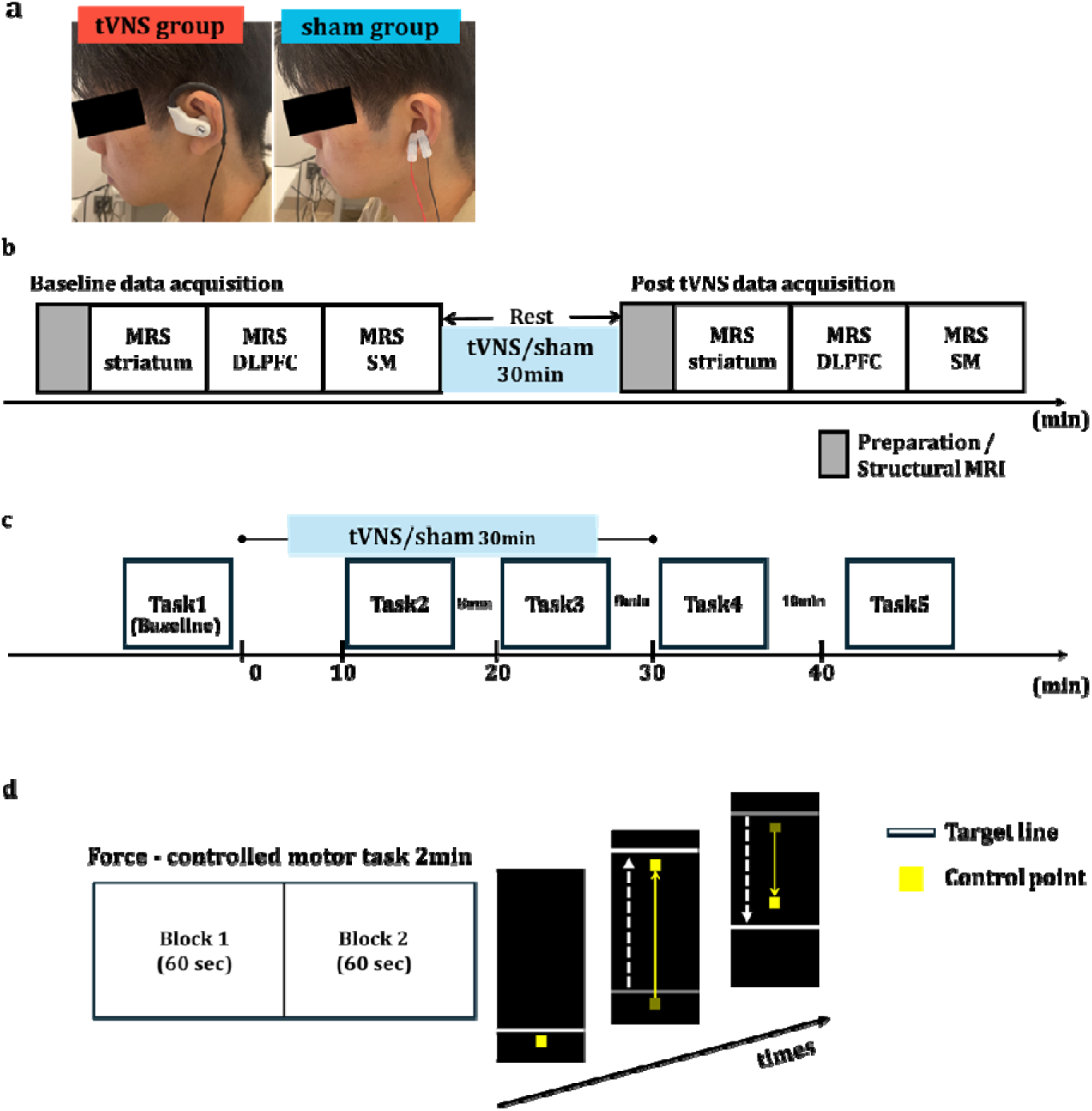
Experimental design **a. Stimulation sites for transcutaneous vagus nerve stimulation (tVNS) and sham groups.** The tVNS group was stimulated at the left cymba conchae (left panel), while the sham group was stimulated at the left earlobe (right panel). **b. Scheme for Experiment 1**. Participants lie in an MRI scanner to measure the baseline GABA concentration in striatum (STR), dorsolateral prefrontal cortex (DLPFC), and sensorimotor cortex (SM). After that, they received either 30 minutes of tVNS or sham stimulation in a separate room. Following stimulation, GABA concentration was remeasured in the same region. **c. Scheme for Experiment 2**. A 2-min force-controlled task was performed before stimulation (Task 1), 10 minutes after stimulation onset (Task 2), 20 minutes after stimulation onset (Task 3), immediately after the end of the stimulation (Task 4), and 12 min after the end of the stimulation (Task 5). **d. Motor task protocol.** Each trial consisted of two 60-s blocks and lasted 2 minutes. Participants were instructed to adjust the pinch force using their right thumb and index finger to quickly and accurately match a randomly moving target line.

Previous studies have shown that a stimulation intensity around 1.0 mA can induced cortical plasticity [47–50], so 1.0 mA was chosen for both tVNS and sham stimulation. To assess the potential side effects of tVNS, all participants completed visual analog scales (VASs) rating pain, itchiness, and discomfort after the experiments.

### 2.3. Experiment 1. Effects of tVNS on regional GABA levels measured by MRS

#### 2.3.1. Participants

Thirty-four healthy adults (17 females, 17 males, mean age = 22.7±2.8 years, range = 19–29 years) were recruited for Experiment 1 to investigate the effects of tVNS on regional brain GABA . Inclusion criteria were (i) right-handed and (ii) 18 to 29 years of age, while exclusion criteria were (i) claustrophobia, (ii) implanted metal objects, (iii) history of psychiatric disorders, and (iv) current use of psychotropic medication. All participants were right-handed as determined by the Edinburgh Handedness Inventory (EHI) [51]. After enrolment, participants were randomly assigned to the tVNS group (n=17) or the sham group (n=17).

#### 2.3.2. Experimental design

After screening for implanted metallic materials, the participants were positioned in the MRI scanner and baseline GABA levels measured using proton magnetic resonance spectroscopy (^1^H-MRS). Following baseline measurements, participants of the tVNS and sham groups received the pre-allocated treatment (tVNS applied to the left cymba conchae or sham stimulation to the left ear lobe for 30 minutes) in a separate room. Post-stimulation GABA levels were then measured during a second MRI session (Figure 1b).

#### 2.3.3. MRS and structural MRI acquisition

Magnetic resonance spectroscopic and structural imaging data were acquired on a 3-T MEGNEATOM Skyra Fit scanner equipped with a 32-channel head coil. All MRS and MRI measurements were performed by a neurosurgeon (author HM) with extensive experience in MRS techniques. T1-weighted images were obtained in the axial, coronal, and sagittal planes to define the STR, DLPFC, and SM as volumes of interest (VOIs). The total volumes for STR, DLPFC, and SM were 30 × 25 × 25 mm^3^, 30 × 25 × 25 mm^3^, and 25 × 25 × 25 mm^3^, respectively. MEGA-PRESS-edited GABA spectra [52,53] were acquired from each VOI using the following parameters: repetition time (TR) = 3 s; echo time (TE) = 68 ms; editing pulse applied either at 3.0 ppm (ON) or 6.4 ppm (OFF); segments of 2048 data points each for ON and OFF acquisitions; spectral bandwidth of 2 kHz interleaved 64 times. In addition, eight dummy pulse scans were performed for signal normalization. In total, 136 scans were acquired from each participant, and total acquisition time was 408 s.

#### 2.3.4. MRS analysis

MRS data were imported into the LCmodel (version 6.3-1N) spectral analysis software program (http://lcmodel.ca/lcmodel.shtml), corrected for frequency and phase drift in 64 edit-ONs and OFFs, and then Gaussian filtered (2 Hz) and Fourier transformed prior to within- and between-group comparisons. GABA+ was obtained from the edited spectra as a ratio relative to NAA using LCmodel. MRS voxels were then coregistered with the structural MRI image of each participant and segmented to determine the fractions of gray matter (GM), white matter (WM), and CSF using GANNET software (version 3.1.5) (https://markmikkelsen.github.io/Gannet-docs/). Regional differences in GABA concentrations are negligible in CSF but approximately twice as large in GM than WM, so tissue correction was applied.

#### 2.3.5. VOI analysis

To ensure that MRS voxels were consistently located in the same brain regions between participants, groups, and pre- and poststimulation scans, voxel masks were normalized to Montreal Neurological Institute space using SPM25 (https://www.fil.ion.ucl.ac.uk/spm/). The spatial overlap between pre- and poststimulation VOIs was then visualized using MRIcroGL (https://www.nitrc.org/projects/mricrogl).

#### 2.3.6. Statistical analysis

Regional GABA levels were compared between tVNS and sham groups by independent samples Student’s t-tests, while the VOI overlap between pre- and post-scans in tVNS and sham groups was compared by paired Student’s t-tests to confirm the consistency of the VOI location.

### 2.4 Experiment 2. Effects of tVNS on motor learning

#### 2.4.1. Participants

Twenty-seven right-handed healthy participants (14 males, 13 females, mean age = 20.9 ± 0.9 years, range = 19–23 years) were recruited for Experiment 2 to investigate the effect of tVNS on motor learning. Inclusion criteria were (i) right-handed and (ii) 18 to 29 years of age, while exclusion criteria were (i) psychiatric diagnoses and (ii) current treatment with psychotropic agents. All participants were right-handed as determined by the EHI (Oldfield, 1971). Again, participants were randomly assigned to a tVNS group (n = 14) or sham group (n = 13). One of the participants recruited in Experiment 1 also participated in Experiment 2. Considering the lasting effects of tVNS, a 4-month interval was maintained for this participant between the two experiments.

#### 2.4.2. Experimental design and motor learning task

In Experiment 2, we investigated the effects of tVNS on motor learning using a time-series design. Participants completed a force-control task at five different time points: before tVNS or sham stimulation (Task 1), 10 min (Task 2) and 20 min (Task 3) after stimulus onset, immediately after stimulation (Task 4), and 12 minutes post-stimulation (Task 5). As in Experiment 1, 30 min of tVNS or sham stimulation was administered during the session, and motor performance was evaluated repeatedly to capture the learning process across time (Figure 1c).

The force-control motor task was conducted as described in a previous study [54]. Briefly, each trial consisted of two, 60-s blocks and lasted 2 minutes. During the task, participants used a pressure sensor (Interlink Electronics Inc., USA) operated with their right thumb and index finger to control the vertical movement of a yellow point on a computer screen (Figure 1d). By adjusting the pinch force, they attempted to align the yellow point with a white target line that moved in a pseudorandom pattern across four force levels (0.5, 1.0, 1.5, and 2.0 N). For each force level, amplitude rose from 0 N to the target value and back to 0 N within 10 s. We calculated the mean absolute error (N) between the target line and the actual force output separately for each block to quantify motor performance (Blocks 1 to 8). To evaluate the effect of stimulation on motor learning, group comparisons were performed at each corresponding block using the unpaired Student’s t-test.

## 3. Results

### 3.1. Experiment 1: Effects of tVNS on regional GABA levels

#### 3.1.1. Side effects of tVNS and sham stimulation

Five of 17 participants reported minor itchiness (mean VAS score, 3.3 ± 2.6) and three minor discomfort (mean 2.3 ± 0.9) during tVNS, but none reported pain. Two of 17 participants in the sham group also reported minor itchiness (mean = 3.4 ± 1.3), whereas none reported pain or discomfort.

#### 3.1.2 Comparisons of regional GABA levels before and after tVNS

No adverse effects of MRS were reported, and GABA-edited MRS data were obtained from all 34 participants in Experiment 1. However, 4 participants (2 in each group) were excluded due to data corruption during acquisition. As a result, data from 15 tVNS group and 15 sham group participants were included in the final analyses (Table 1, upper). Before analyzing GABA levels, we compared the VOI overlap for each target region between treatment groups, and found no significant differences (Figure 2a, Table 2). Thus, differences in GABA were not influenced by inaccuracies in setting VOI boundaries.

**Figure 2.**
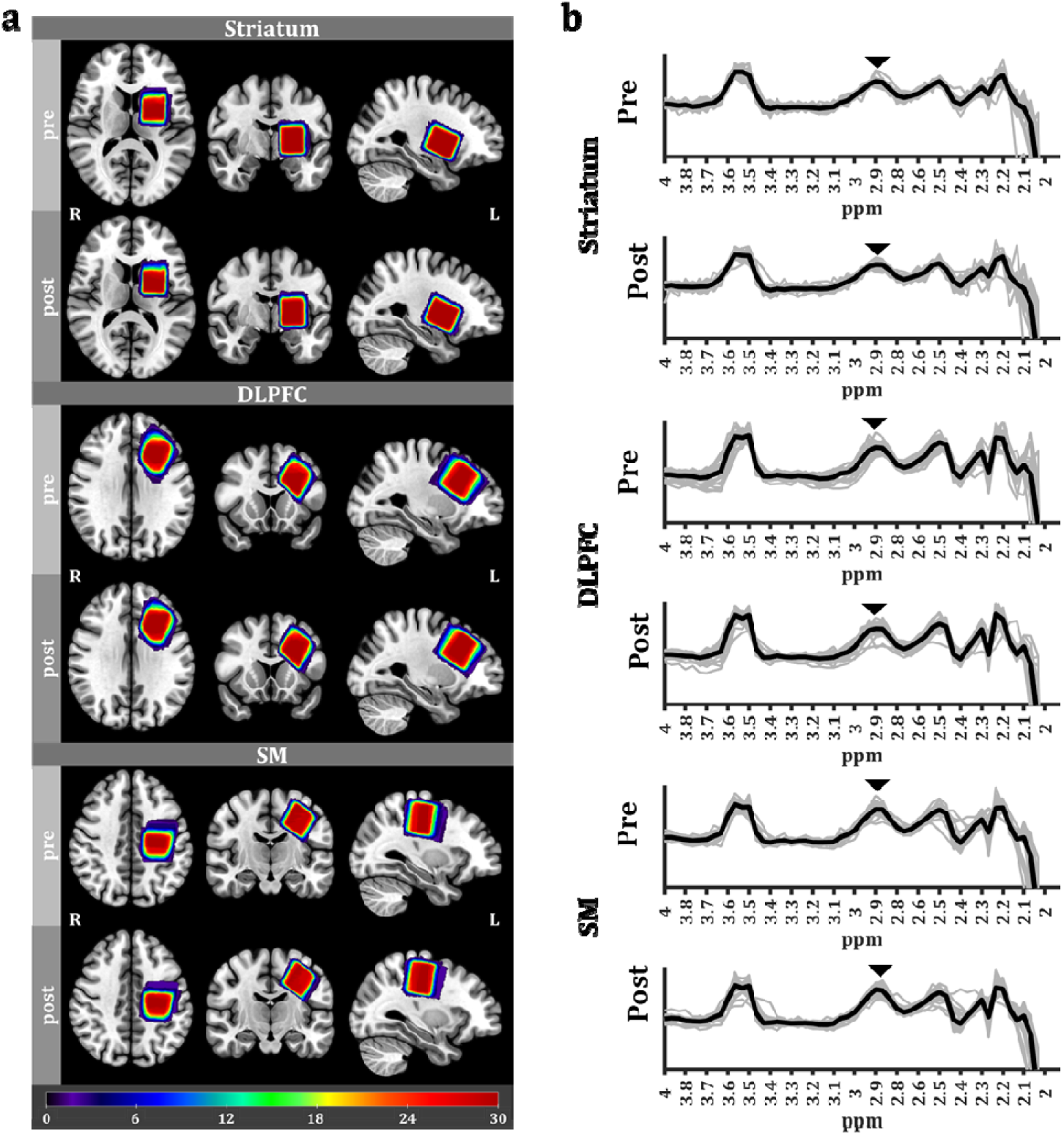
VOI analysis and the MRS-GABA spectrum **a. VOI analysis.** Heatmaps representing the spatial overlap across participants for STR, DLPFC, and SM MRS voxels (pre; upper, post: bottom). Color bars represent the number of overlapping voxels. Heatmaps are overlaid over the mean structural image across all participants. The high degree of spatial overlap indicates a high consistency in voxel placement across time points and individuals. **b. Pooled pre- and poststimulation MRS-GABA spectrum data in each brain region.** Black solid lines indicate averaged MRS spectrum data across all participants, and gray solid lines indicate individual-participant MRS spectrum data. Black arrowheads indicate the GABA peaks.

**Table 1.**
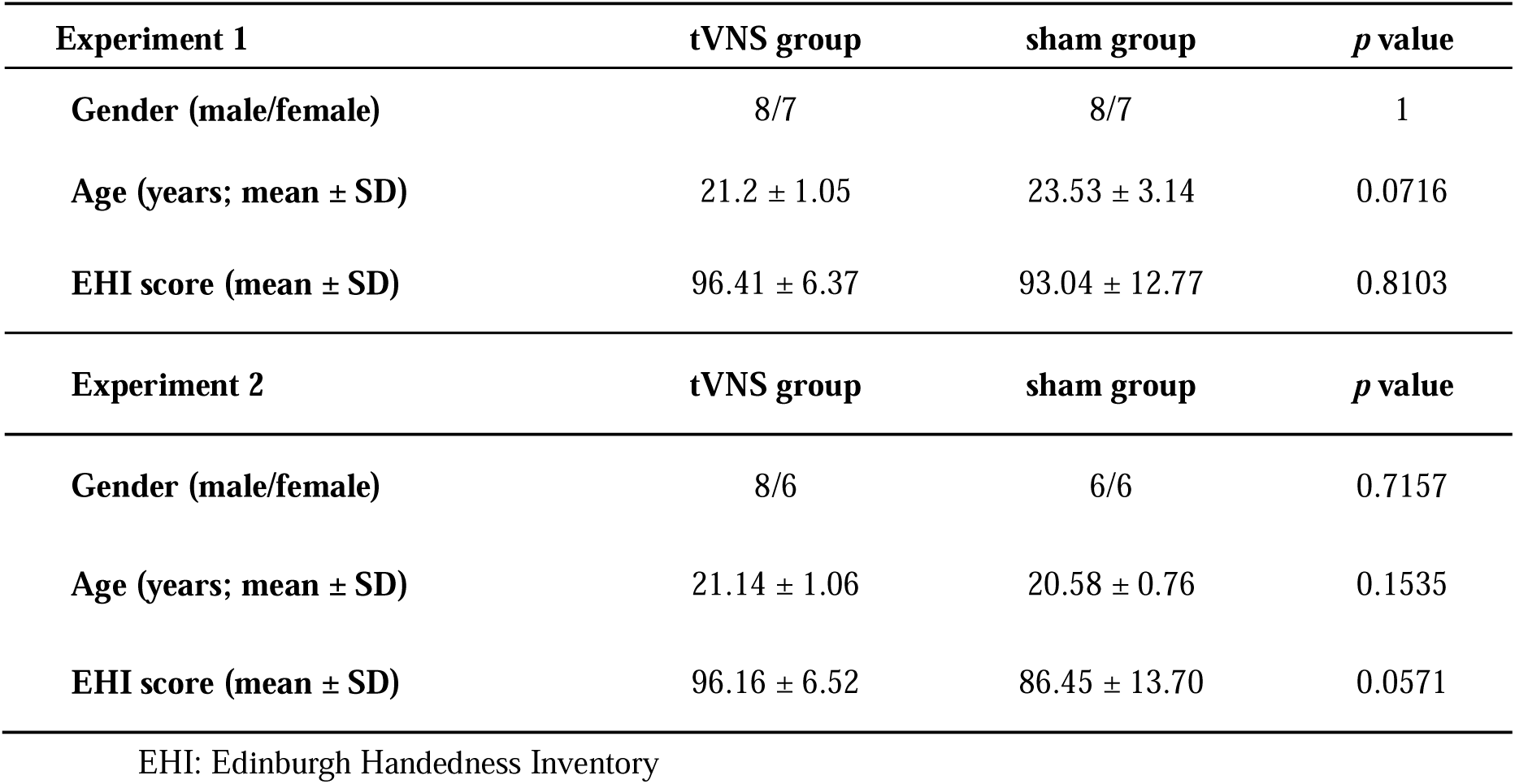
Demographic characteristics of the participants.

**Table 2.**
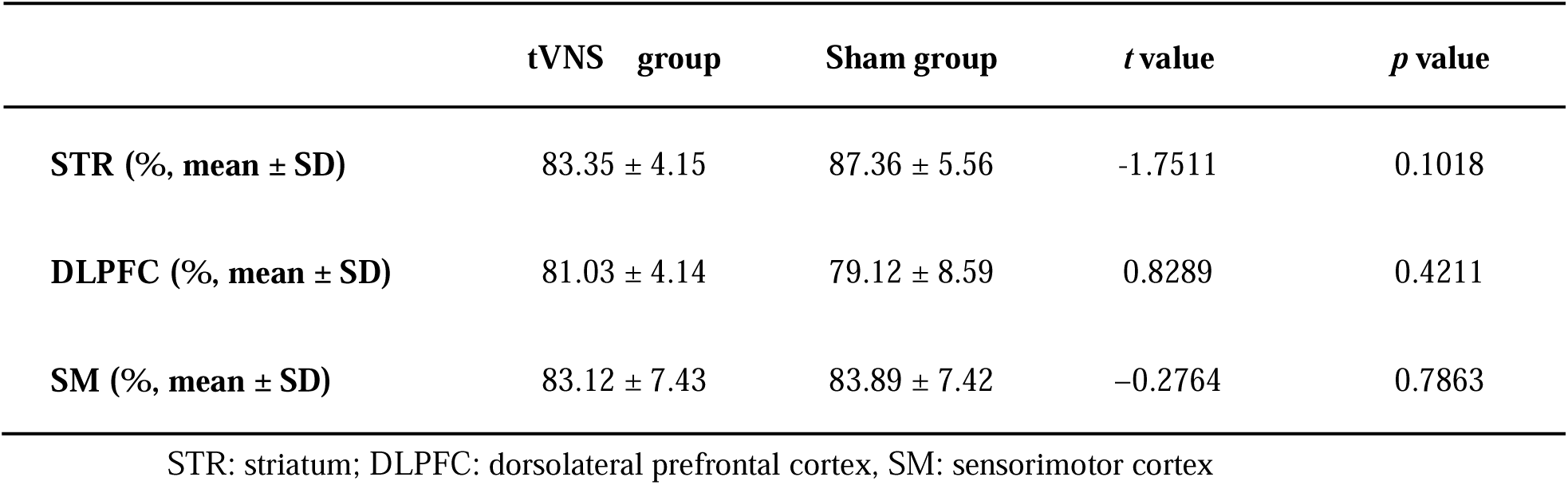
Regional volume of interest overlap between measurement times.

Regional MRS-GABA spectra following tVNS or sham stimulation were then compared to corresponding baselines (Figure 2b). Striatal (STR) total GABA level was significantly lower following tVNS compared to baseline (*p* = 0.041, *t-value* = 1.861, *Cohen’s d* = 0.481) (Figure 3), while no significant changes were observed in the DLPFC (*p* = 0.222, *t-value* = 0.786, *Cohen’s d* = 0.203) and SM (*p* = 0.465, *t-value* = 0.090), *Cohen’s d* = 0.023), although there was a numeric decrease in the DLPFC. Alternatively, no significant changes were observed in the STR (*p* = 0.180, *t-value* = 0.943, *Cohen’s d* = 0.066), DLPFC (*p* = 0.890, *t-value* = −1.286, *Cohen’s d* = 0.325), and SM (*p* = 0.562, *t-value* = −0.160, *Cohen’s d* = −0.058) following sham stimulation, although a numeric increase was observed in the DLPFC.

**Figure 3.**
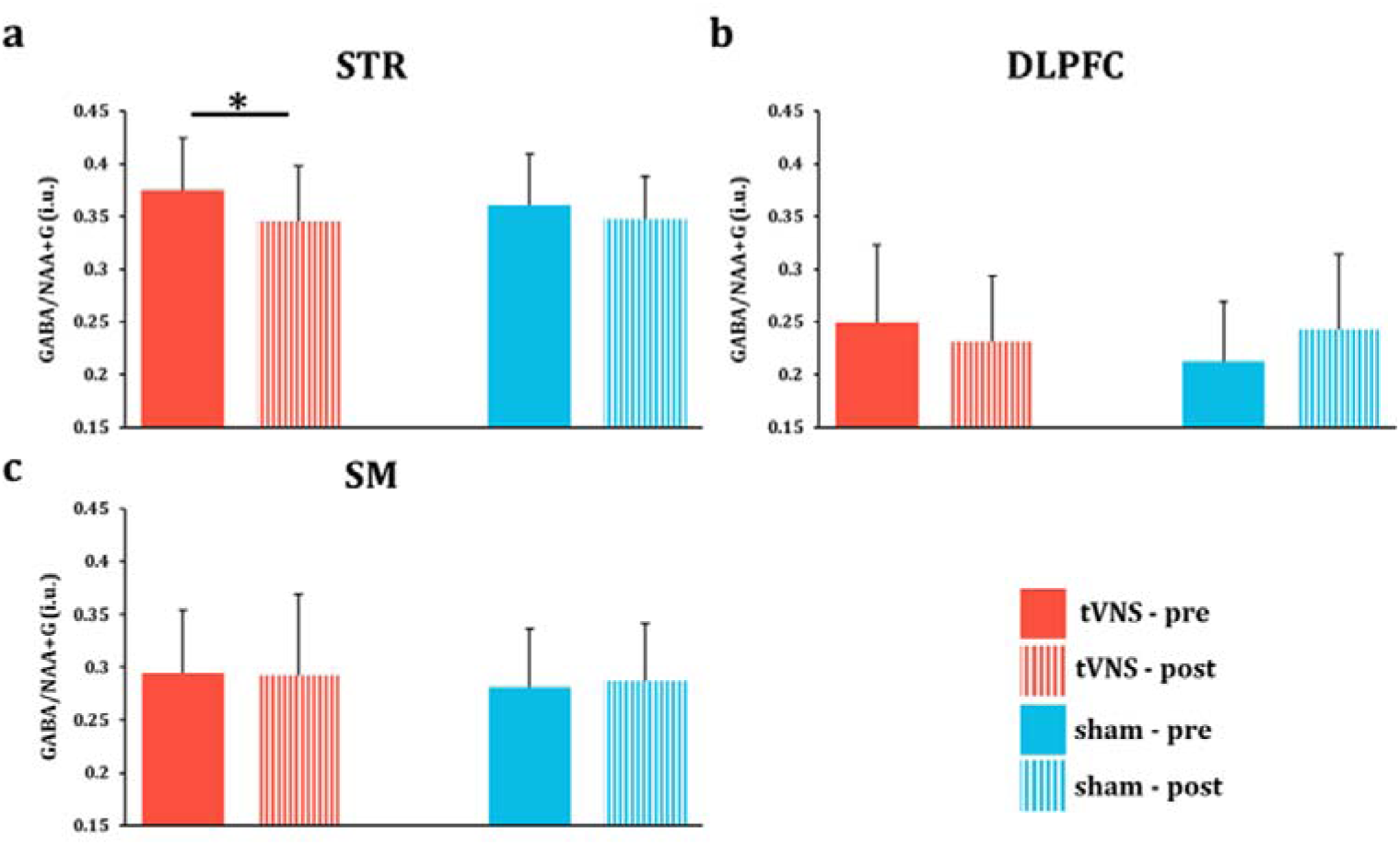
tVNS reduced total GABA concentration in the striatum **a.** Striatal GABA was significantly reduced following tVNS but not sham stimulation (* *p* < 0.05). No significant changes were observed in the DLPFC or SM. STR: striatum, DLPFC: dorsolateral prefrontal cortex, SM: sensorimotor cortex. Error bars indicate the SD.

### 3.2. Experiment 2: Effects of tVNS on motor task performance

#### 3.2.1 Side effects

Two participants reported mild pain (mean VAS score 2.6 ± 1.0), four reported moderate itchiness (mean 4.3 ± 1.5), and four experienced minor discomfort (mean 1.5 ± 0.8) following tVNS. Two participants reported mild pain (mean 2.3 ± 0.8), one reported itchiness (1.6), and one felt discomfort (2.3) following sham stimulation.

#### 3.2.2 Task performance improvement by tVNS

One participant who could not adequately manipulate the force sensor in the force-control motor task was excluded from the analysis. Thus, the final sample sizes were 14 for the tVNS group and 12 for the sham group (Table 1, lower). The mean error values in each block were defined as learning values and compared across blocks. Mean error values were significantly lower following 10 min of tVNS compared to 10 min of sham stimulation (*p* = 0.0232, *t-value* =−2.635, *Cohen’s d* = −1.0367). Alternatively, performance levels did not differ after 20 min of tVNS, immediately after 30 min of tVNS, or 12 min after 30 min of tVNS compared to the sham stimulation condition.

## 4. Discussion

### 4.1. Effects of tVNS on GABA levels

In Experiment 1, we found that tVNS reduced total GABA content in the STR, although no significant changes were observed in the DLPFC or SM. A previous study [55] similarly reported that tVNS reduced GABA levels in the anterior cingulate cortex of a patient with treatment-resistant depression. Thus, tVNS may not simply enhance regional GABA contents as reported in multiple previous studies, but may also reduce levels, at least in some regions, an effect that could in turn facilitate neuroplasticity through disinhibition of various motor-related circuits. Indeed, we observed enhanced learning of a force-control motor task during tVNS.

While this effect appeared early and transient (only after 10 min of stimulation), task performance was also rapidly reaching a plateau. Therefore, additional studies are needed to examine if tVNS enhances motor learning across phases and task conditions, such as during more difficult tasks with no rapid ceiling effect (or floor effect on error rate as measured in the present study). If so, tVNS could be a broadly useful adjunct treatment for recovery of motor function.

The striatum contains abundant GABAergic interneurons and short projection neurons that facilitate motor learning by modulating cortical plasticity through direct and indirect pathways [56]. The present findings suggest that a decrease in striatal GABA and ensuing disinhibition may have contributed to faster learning of the force-control task via effects on corticostriatal plasticity. Frrand et al. [57] reported that iVNS increased the expression of multiple neurotransmitters in the STR and concomitantly mitigated motor deficits in a rat model of Parkinson’s disease, possibly by enhancing striatal output through disinhibition and strengthening of network connectivity as a previous reported that tVNS enhanced functional connectivity within the striatum-prefrontal cortex network [58]. Additionally, tVNS increased BOLD signals in the STR [33] and there is a negative correlation between GABA levels and BOLD signals [28–32], further supporting a tVNS-induced decrease in striatal GABA levels.

Although no significant differences were observed, tVNS also numerically reduced GABA in the DLPFC, whereas the sham group showed an increase, suggesting enhanced activation of STR–DLPFC circuits by tVNS. Capone et al. [59] reported that left-sided tVNS increased short-interval intracortical inhibition, which is reflective of GABA-A inhibitory circuit activity, in the right SM but not the left SM, so enhanced GABAergic inhibitory activity in the right (or contralateral) SM cannot be ruled out. Alternatively, it is also possible that the absence of observable GABA modulation in the SM may be due to the delay in MRS acquisition (∼40 minutes after tVNS).

### 4.2. Acceleration of motor learning by tVNS

Administration of tVNS for 10 min enhanced motor learning, although not maximum performance, suggesting a specific influence on early-phase motor learning. However, as discussed, longer term effects during stimulation may have been obscured by floor effect on the error rate, so further studies are warranted to examine the efficacy for improving performance on more difficult motor tasks.

Suppression of GABA receptor functions has been reported to enhance motor learning consolidation and motor memory reconsolidation [60]. Similarly, Stagg et al. concluded that a reduction in GABAergic activity can enhance motor learning, while Cardellicchio et al. reported that high GABA levels in the sensorimotor cortex inhibited motor learning [61]. In accord with this reciprocal relationship between striatal GABAergic activity and motor learning, tVNS both enhanced early-phase motor learning and reduced striatal GABA. A previous animal study identified the posterior dorsomedial striatum as a critical area for motor learning by associating sensory information with action outcomes [62]. Human studies [63,64] have also suggested that the STR is essential for early-phase motor learning. Cataldi et al. [65] and Yin et al. [66] reported that the caudate nucleus, a component of the STR, is involved in the early, rapid phase of motor learning while others have implicated the putamen (the other STR component) in later stages of learning. A functional MRI study [67] also revealed enhanced activity in the head of the caudate nucleus during the initial learning phase. Furthermore, recent studies have reported that noninvasive brain stimulation targeting the STR, such as transcranial temporal interference stimulation, can enhance striatal activity and facilitate motor learning [68–70]. Given these similarities, it is conceivable that tVNS may serve as an alternative noninvasive neuromodulation approach to activate (disinhibit) the striatum and promote motor learning.

Although tVNS promotes early-phase motor learning, we cannot rule out the possibility that tVNS may also enhance the overall capacity for motor skill acquisition. In our study, performance improvements reached a plateau around Task 3, approximately 20 minutes after tVNS initiation (Figure 4). This ceiling effect may have limited our ability to detect further enhancements in motor performance. Employing a task with higher difficulty or extended duration may allow for a more accurate assessment of the potential effects of tVNS on the full capacity of motor learning. Future studies are warranted to explore this possibility.

**Figure 4.**
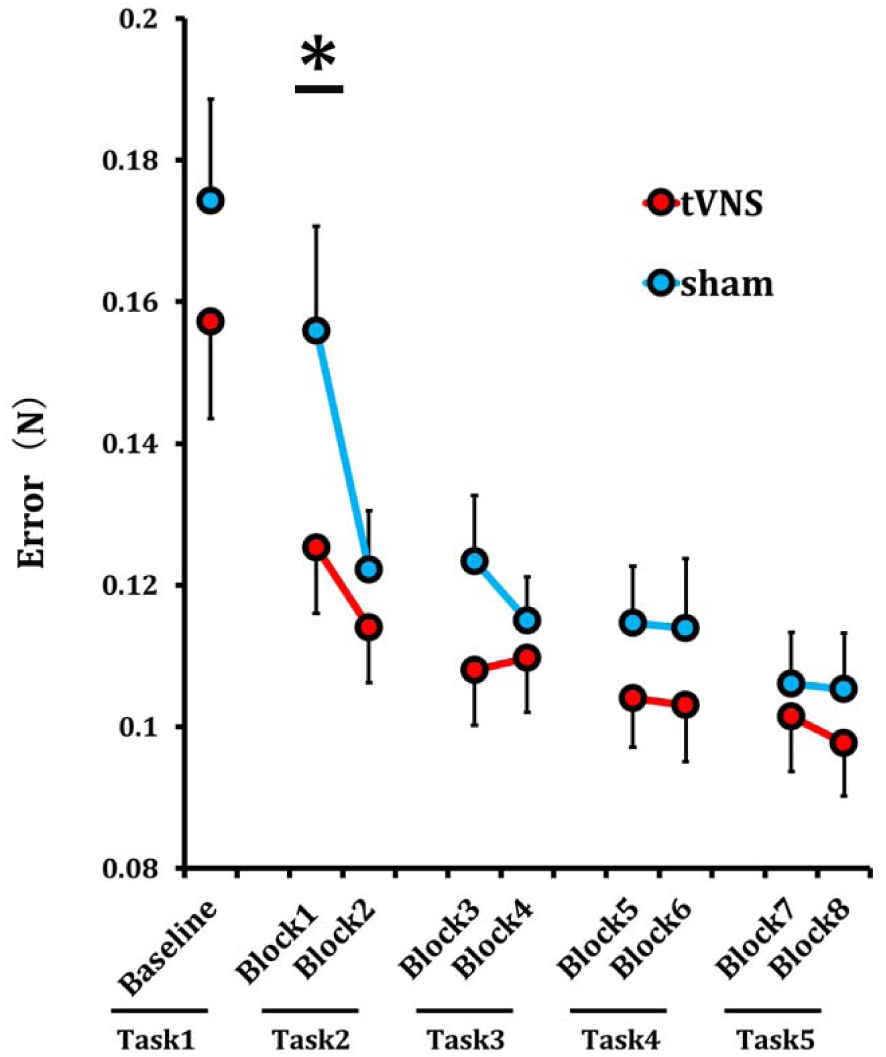
tVNS improved force-control motor task performance Time course of the mean error during the force-control motor task. The red line represents the tVNS group and the blue line represents the sham group. Each task consisted of two blocks. Error values were averaged within each block and corresponding blocks compared between tVNS and sham groups. In block 1 (10 minutes post-stimulation), the tVNS group showed a significantly lower error value compared with the sham group (\**p* < 0.05). Error bars indicate the SD.

### 4.3. Clinical implication

The observed reduction in striatal GABA levels and acceleration of early-phase motor learning suggest that tVNS may be a useful intervention for post-stroke rehabilitation as excessive tonic inhibition has been shown to hamper motor recovery. As a noninvasive and portable technique, tVNS holds practical advantages over other neuromodulation methods and may be more easily implemented in rehabilitation settings. Further research is needed to determine its efficacy in patient populations and to optimize stimulation protocols for clinical use.

### 4.4. Limitations

Our study has several limitations. First, Experiments 1 and 2 were conducted separately in different participant groups. Therefore, we could not establish a direct correlation between changes in striatal GABA content and motor learning performance. Second, it is technically challenging to measure GABA levels in multiple brain regions simultaneously using MRS. As a result, there was a time lag between scans for different brain regions, and transient changes induced by tVNS may have been missed. In addition, while MRS offers the advantage of non-invasive measurement, it does not distinguish between intracellular and extracellular GABA and thus among changes in synthesis, storage, and release into synaptic and presynaptic spaces. Third, while a previous study [71] reported a lasting effect of tVNS on depression scores, our study did not examine the after-effect of tVNS on GABAergic modulation and motor learning.

## CRediT authorship contribution statement

**Kana Matsumura:** Writing–original draft, Investigation, Formal analysis, Data curation, Visualization **Hiroyuki Matsuta:** Writing–original draft, Formal analysis, Methodology, Data curation. **Ryushin Kawasoe:** Data curation. **Tomoyuki Fumuro:** Data curation. **Kojiro Matsushita;** Software **Nobuhiro Hata;** Investigation, Resources. **Yoshiki Asayama;** Investigation, Resources **Tsuyoshi Shimomura;** Investigation, Resources **Minoru Fujiki:** Investigation, Resources, and Supervision **Hisato Sugata:** Project administration, Writing–review & editing, Validation, Supervision, Methodology, Investigation, Funding acquisition, Data curation, Conceptualization.

## Data availability

Anonymized data supporting the conclusions of this study can be obtained from the corresponding authors on reasonable request.

## Acknowledgments

We thank all participants in this study and Enago (www.enago. com) for the English language review.

